# Broad-spectrum antiviral activity of the synthetic rocaglate zotatifin against multiple viruses

**DOI:** 10.1101/2025.11.18.689102

**Authors:** Patricia Valdés-Torres, Dalkiria Campos, Paola Elaine Galán-Jurado, Dalel Zegarra, Isaac Tuñon-Lorenzo, Félix González-Castillo, María Blanquer, Juan Castillo Mewa, Carmen Rivas, José González-Santamaría

## Abstract

Viruses pose a significant threat to global public health, yet therapeutic options are limited. Zotatifin, a synthetic rocaglate, targets eukaryotic initiation factor 4A (eIF4A), thereby inhibiting viral protein synthesis and triggering an interferon response. Whether zotatifin exhibits broad-spectrum antiviral activity is unknown. Here we evaluated zotatifin’s cytotoxicity using the MTT method and its antiviral activity using plaque-forming assays, Western blot analysis, immunofluorescence, and flow cytometry across multiple cell lines and virus strains. Zotatifin demonstrated better tolerability than rocaglates rocaglamide A and CR-1-31-B. The compound exhibited potent, dose-dependent inhibition of the Mayaro virus, achieving greater than a 4-log reduction in viral titers at a 50 nM dose, regardless of the cell line or virus strain tested. In addition, zotatifin was effective to inhibit multiple arboviruses (Chikungunya, Una, and Zika), influenza A virus, vesicular stomatitis virus, and vaccinia virus. Zotatifin down-regulated viral protein synthesis for all viruses tested and analysis of gene expression using RT-qPCR revealed activation of the type I interferon pathway in zotatifin-treated cells. Taken together, these results suggest that zotatifin exhibits broad-spectrum antiviral activity against five virus families, likely via several molecular mechanisms. This work supports the potential therapeutic use of zotatifin as a pan-antiviral drug in humans.

## 1. Introduction

Despite the considerable risks that viruses pose to human health, effective antiviral drugs and licensed vaccines remain unavailable for many of these pathogens. This therapeutic gap has prompted researchers to explore novel antiviral strategies, particularly those targeting host factors essential for viral replication. Recently, rocaglates, a class of naturally occurring compounds derived from *Aglaia* plants, have emerged as promising broad-spectrum antiviral agents (Agarwal et al., 2021; Harneti & Supratman, 2021).

Rocaglates exert antiviral properties primarily through interaction with eukaryotic initiation factor 4A (eIF4A), a DEAD-box RNA helicase and key component of the protein synthesis machinery (Greger, 2022; Li et al., 2023; Li-Weber, 2015). eIF4A is essential for the translation of structured mRNAs, including many viral transcripts that contain complex 5’ untranslated regions (UTRs) (Taroncher-Oldenburg et al., 2021). By targeting eIF4A, rocaglates effectively block viral protein synthesis, thereby inhibiting viral replication while showing selectivity for viral over cellular mRNAs (Taroncher-Oldenburg et al., 2021).

The natural rocaglate silvestrol has demonstrated remarkable antiviral activity against multiple viral pathogens, including Middle East respiratory syndrome coronavirus (MERS-CoV), human coronavirus 229E (HCoV-229E), influenza A virus (IAV), Ebola virus (EBOV), hepatitis E virus, and Zika virus (ZIKV) (Biedenkopf et al., 2017; Elgner et al., 2018; Glitscher et al., 2018; Muller et al., 2018; Slaine et al., 2017). Building on these findings, researchers have developed synthetic rocaglates with comparable efficacy. For instance, the synthetic compound CR-1-31-B has been reported to exhibit antiviral activity like silvestrol (Müller et al., 2020).

More recently, zotatifin (eFT226), another synthetic rocaglate (Ernst et al., 2020), has shown particularly promising results. This compound effectively controls diverse coronaviruses replication and reduces SARS-CoV-2 viral infectivity (Gordon et al., 2020; Obermann et al., 2022). Importantly, zotatifin exhibits reduced cytotoxicity toward immune cells compared to other rocaglates and has entered clinical trials for cancer (ClinicalTrials.gov: NCT04092673) and COVID-19 (ClinicalTrials.gov: NCT04632381) (Obermann et al., 2022). Recent studies have revealed that zotatifin may have other mechanisms of action beyond eIF4A inhibition such as the induction of the interferon (IFN) pathway, which may contribute to its antiviral efficacy (Zhao et al., 2023). However, despite these encouraging findings, zotatifin’s antiviral spectrum against viral families beyond coronaviruses remains largely unexplored.

In this study we have evaluated zotatifin’s potential to control viral replication across diverse viral families, including representatives from the *Togaviridae* (Mayaro, Chikungunya, and Una viruses), *Flaviviridae* (Zika virus), *Orthomyxoviridae* (influenza A virus), *Rhabdoviridae* (vesicular stomatitis virus), and *Poxviridae* (vaccinia virus). Our results demonstrated that zotatifin exhibits broad-spectrum antiviral activity against five virus families, supporting its potential therapeutic use as a pan-antiviral drug in humans.

## 2. Materials and Methods

### 2.1 Cell lines, virus strains, and reagents

Human dermal fibroblasts (HDFs, Cat. # PCS-201-012), human immortalized microglial cells (HMC3, Cat. # CRL-3304), human non-small cell lung cancer A549 cells (CCL-185), BSC40 (Cat. # CRL-2761), and Vero-E6 cells (CRL-1586) were obtained from the American Type Culture Collection (ATCC, Manassas, VA, USA). HDFs and A549 cells were cultivated in Dulbecco’s Modified Eagle Medium (DMEM), HMC3 cells in Eagle’s Minimum Essential Medium (EMEM), and Vero-E6 in Minimal Essential Medium (MEM). All media were supplemented with 10% fetal bovine serum (FBS), 2 mM L-glutamine, and 1% penicillin-streptomycin solution. Cells were maintained at 37°C in a humidified atmosphere with 5% CO_2_.

The Mayaro virus (MAYV) strains AVR0565 (Peru), TRVL 4675 (Trinidad and Tobago), BeH256 (Brazil), and D218 (Suriname), as well as Una virus (UNAV) BT 1495-3 (Panama) were obtained from the World Reference Center for Emerging Viruses and Arboviruses (WRCEVA) at the University of Texas Medical Branch (UTMB) in Galveston, Texas, USA and were kindly provided by Dr. Scott Weaver. Chikungunya (CHIKV, Panama_256137) and Zika (ZIKV, 259249) viruses were isolated from patients in Panama. The recombinant GFP-expressing vesicular stomatitis virus (rVSV-GFP) and influenza A virus (PR8-GFP) were provided by Dr. Adolfo García-Sastre (Icahn School of Medicine at Mount Sinai in New York, USA). Vaccinia virus (VV) WR strain was provided by Dr. Mariano Esteban (Centro Nacional de Biotecnología, CSIC, Spain). All virus strains were propagated in Vero-E6 or A549 cells, titrated, aliquoted, and stored at -80°C as previously reported (Valdes-Torres et al., 2022). The rocaglates rocaglamide A (Cat. # 19356), CR1-31-B (Cat. # 136453), and zotatifin (also known as eFT226, Cat. # HY-112163) were purchased from MedChemExpress (New Jersey, USA). All compounds were dissolved in DMSO and stored at -20 °C until use. Working solutions were prepared in appropriate culture media at indicated concentrations immediately before use.

### 2.2 Cell viability assay

Cytotoxicity of rocaglates was evaluated using the MTT assay as previously described (Valdes-Torres et al., 2022). Briefly, cells were seeded in 96-well plates at density of 1x10^4^ cells per well and allowed to adhere overnight. Cells were then treated with different concentrations of rocaglamide A, CR1-31-B, or zotatifin. After 24 hours of incubation, cells were treated with 20 μL of 5 mg/ml solution of 3-(4,5-dimethyl-2-thiazolyl)-2,5-diphenyl tetrazolium bromide (MTT, Sigma-Aldrich, St. Louis, Missouri, USA) and incubated for an additional 2 hours. The formazan crystals were then dissolved in 100 μL DMSO, and absorbance at 570 nm was measured using a microplate reader (BioTek, Winooski, VT, USA). The results are presented as the percentage of viable cells relative to DMSO-treated control cells.

### 2.3 Plaque-forming assay

Vaccinia virus was harvested from infected cells by three cycles of freezing/thawing, sonication and clarification by centrifugation. Clarified supernatants and virus titers in cell culture supernatants were quantified using plaque-forming assays as previously reported (Valdes-Torres et al., 2022). Briefly, 10-fold serial dilutions of infected samples were used to infect confluent Vero-E6, A549 or BSC40 cells in 12-well plates. After 1 hour of incubation at 37 °C, supernatants were removed, and the cells were overlaid with a solution of 1% agar in MEM supplemented with 2% FBS. Plates were incubated at 37 °C until plaques were visible (24-72 hours depending on the virus). The agar overlay was removed, and the cells were fixed with 4% formaldehyde in PBS for 20 minutes, then stained with 2% crystal violet in 30% methanol. Plaques were counted manually, and the virus titers were expressed as plaque-forming units per milliliter (PFU/ml).

### 2.4 Virus infection and cytopathic effect assays

HDFs, HMC3, and A549 cells were seeded in 12- or 24-well plates at appropriate densities and allowed to adhere overnight. Cells were pretreated with DMSO (vehicle control), rocaglamide A, CR-1-31-B, or zotatifin at indicated concentrations for 2 hours at 37°C. Following pretreatment, cells were infected with MAYV, CHIKV, UNAV, ZIKV, PR8-GFP, or rVSV-GFP at specified multiplicities of infection (MOI). After virus adsorption, the inoculum was removed and fresh medium containing the respective compounds was added. At the indicated hours post infection, cell culture supernatants were collected for virus titration, and cells were processed for protein analysis. When the experiments were carried out with VV, cells were collected to be used for either virus titration of for protein analysis, as indicated.

For cytopathic effect (CPE) analysis, cell morphology was evaluated under an inverted microscope at 48 hours post-infection, and representative images were captured (Valdes-Torres et al., 2022).

For time-of-addition studies, the following protocols were used:

- Pretreatment assay: cells were treated with zotatifin for 2 hours, then washed with PBS and infected with virus in compound-free medium.
- Binding assay: cells were cooled at 4°C, incubated with virus and treated simultaneously with zotatifin for 1 hour. Following incubation, cells were washed with PBS and incubated in compound-free medium.
- Entry assay: cells were infected at 4°C for 1 hour, transferred to 37°C, and zotatifin was added for 2 hours. Following incubation, cells were washed with PBS and incubated in compound-free medium.
- Post-entry assay: zotatifin was added at 2 hours post-infection at 37°C and maintained until harvest at 24 hours.

In all assays, viral titers were quantified at 24 hours post-infection.

### 2.5 Flow cytometry assay

A549 cells were seeded in 6-well plates and treated with DMSO or zotatifin (50 nM) for 2 hours, then infected with PR8-GFP or rVSV-GFP at MOI 0.1 or 1.0 PFU/cell and then incubated in the presence or absence of zotatifin. Twnty-four hours later, the cells were harvested by tripsinization, washed with PBS, and fixed with 2% paraformaldehyde for 20 minutes at room temperature. Fixed cells were washed twice and resuspended in 250 μL PBS. Samples were analyzed using a CytoFLEX S flow cytometer (Beckman Coulter, Florida, USA) by acquiring at least 10,000 events per sample. GFP fluorescence was detected using 488-nm excitation and 525-540 nm emission filters. Data were analyzed using CytoExpert software (v2.4.0.28) with appropriate compensation controls.

### 2.6 Western blot assay

Protein extracts were obtained from mock- or virus-infected cells treated or untreated with zotatifin, using Laemmli buffer containing 10% dithiothreitol (Bio-Rad, Hercules, CA, USA). The proteins were then separated using SDS-PAGE on 10-12% Bis-Tris gels, transferred to nitrocellulose membranes. Membranes were blocked with 5% nonfat milk in T-TBS buffer for 30 minutes at room temperature, then incubated overnight at 4 °C with primary antibodies: rabbit polyclonal anti-E1 (1:1000) and rabbit polyclonal anti-nsP1 (1:1000) for MAYV, UNAV, and CHIKV (Llamas-González et al., 2019); anti-nsP2 of CHIVK (Cat. # GTX135188, GeneTex, USA); anti-NP of IAV (1:1000) (GTX125989, USA); and anti-GAPDH (1:1000, Cat. # VMA00046, Bio-Rad); or anti-β-actin (1:1000, Cat. # VMA00048, Bio-Rad). After washing three times with T-TBS, membranes were incubated with HRP-conjugated goat anti-rabbit (Cat. # 926-80011, LI-COR, Lincoln, NE, USA) or goat anti-mouse (Cat. # 926-80010, LI-COR, Lincoln, NE, USA) secondary antibodies for 1 hour at room temperature. Chemiluminescent signals were detected using SignalFire^TM^ ECL Reagent (Cell Signaling Technology, Danvers, MA, USA) and visualized using a C-Digit scanner (LICOR, Lincoln, NE, USA) or a film system.

### 2.7 Immunofluorescence assay

A549 cells grown on glass coverslips in 24-well plates were treated with or without zotatifin (50 nM) for 2 hours, then infected with the MAYV ARV0565 strain at a MOI of 1 in presence or absence of the compound. At 24 hours post infection, cells were fixed with 2% paraformaldehyde for 20 minutes, permeabilized with Triton X-100 for 10 minutes, and blocked with 5% bovine serum albumin as previously described (Sugasti-Salazar et al., 2021). Cells were stained with rabbit anti-E1 or anti-nsP1 primary antibodies (1:100) overnight at 4°C, followed by incubation with Alexa Fluor 568 goat anti-rabbit secondary antibody (1:1000, Invitrogen, Carlsbad, USA) for 1 hour at room temperature. Coverslips were mounted using Prolong Diamond Antifade with DAPI (Invitrogen, Carlsbad, USA). Images were acquired using an FV3000 Fluoview confocal microscope (Olympus, USA).

### 2.8 Gene expression analysis by RT-qPCR

Total RNA was extracted from A549 cells treated with or not with zotatifin (50 nM) for 8 hours using the RNeasy kit (QIAGEN, Valencia, CA, USA), according to the manufactureŕs instructions. Single-stranded cDNA was synthesized from 1 μg of RNA using a High-Capacity cDNA Reverse Transcription kit. RT-qPCR was performed using Power SYBR Green PCR Master Mix in a QuantiStudio 5 thermocycler (Applied Biosystems, Foster City, CA, USA) and the primers indicated in Table 1, as previously described (Valdes-Torres et al., 2022). Relative mRNA expression was calculated using the β-actin gene for normalization with the ΔΔCT method (Livak & Schmittgen, 2001).

**Table 1.**
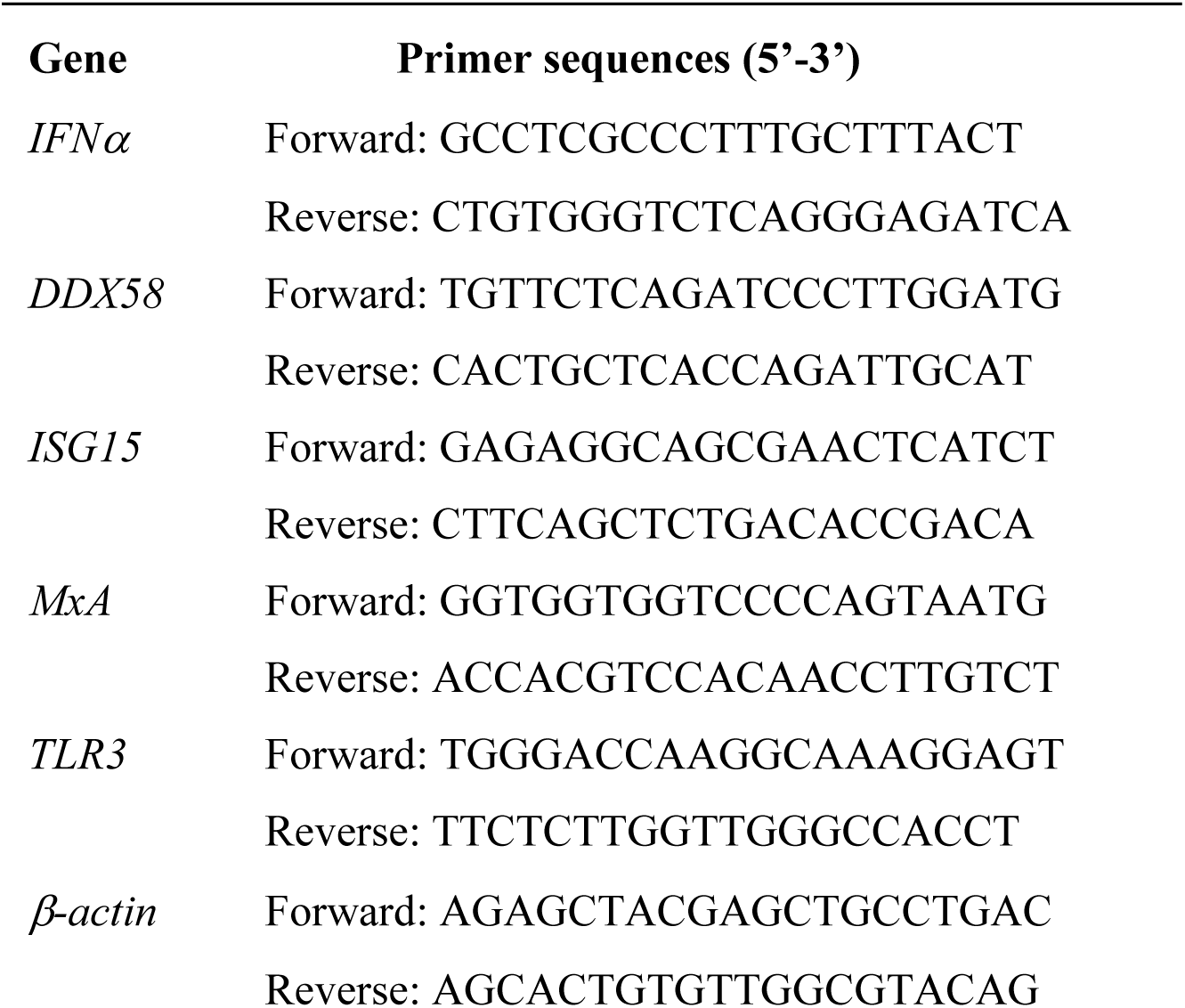
Primers used in this study.

### 2.9 Data analysis

All experiments were performed in biological triplicate. Data are present as mean ± standard deviation. Statistical comparisons were performed using one-way ANOVA followed by Dunnettt’s post hoc test for multiple comparisons against control, or unpaired t-test for pairwise comparisons. A p-value < 0.05 was considered statistically significant. All statistical analysis and graphics were generated using GraphPad Prism software (version 10.6.0 for Mac; GraphPad Software, San Diego, CA, USA).

## 3. Results

### 3.1 Zotatifin potently controls Mayaro virus replication

First, we compared zotatifin cytotoxicity profile with those of other rocaglates, including the natural compound rocaglamide A and the synthetic CR-1-31-B (Figure 1A). Using the MTT assay, we assessed cytotoxicity in primary human dermal fibroblasts (HDFs) and human immortalized microglia cells (HMC3). Rocaglamide A and CR-1-31-B exhibited notable toxicity at doses exceeding 5 nM in both cell lines (Figure 1B and C; <80% cell viability). In contrast, zotatifin demonstrated favorable tolerability in these human cell lines at higher doses (≥80% cell viability at 50 nM Zotatifin) (Figure 1D).

**Figure 1.**
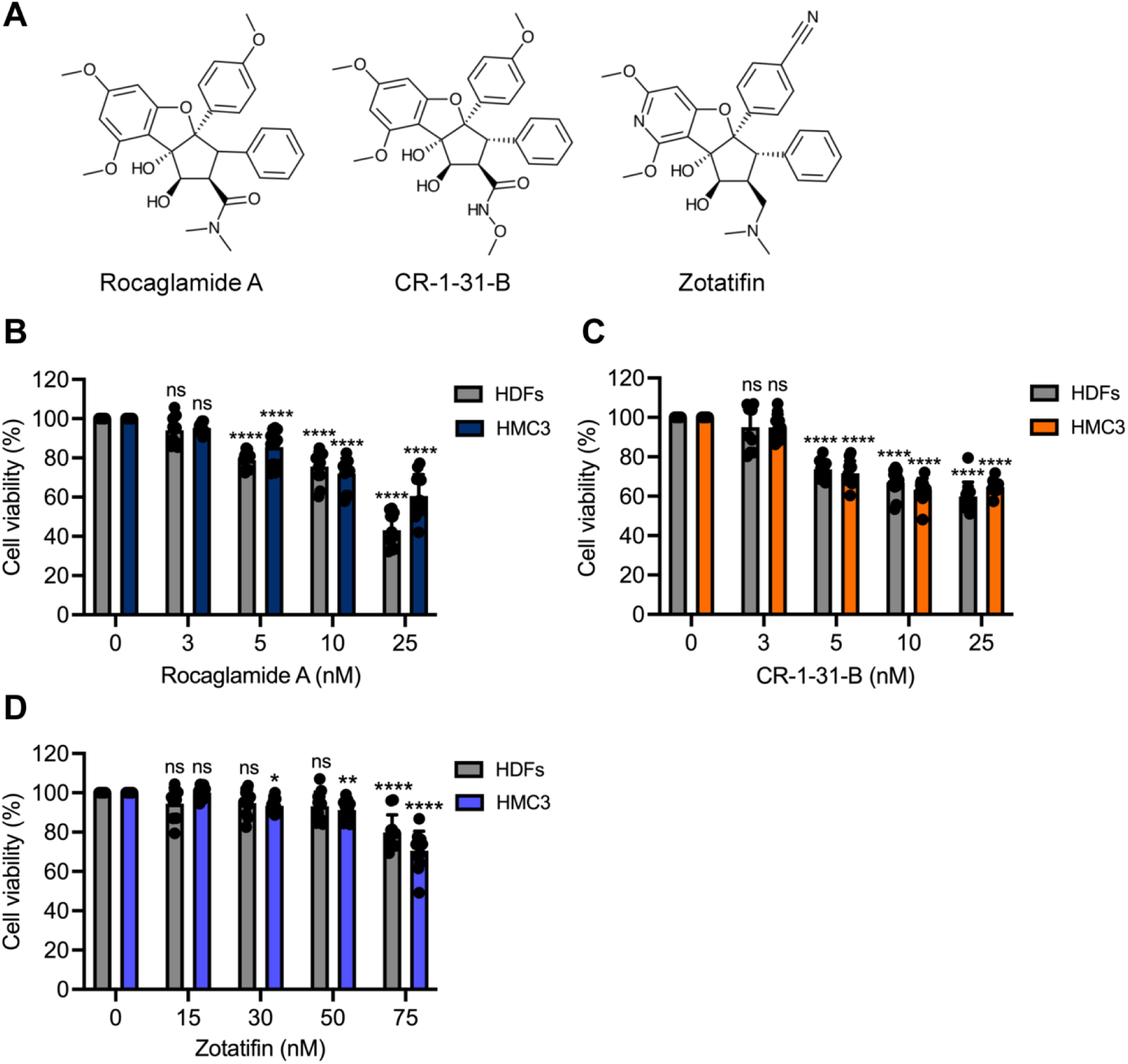
Cytotoxicity of rocaglates in human dermal fibroblasts (HDFs) and microglial cells (HMC3). (**A**) Chemical structures of the rocaglates tested in this study. HDFs and HMC3 cells were treated with increasing concentrations of rocaglamide A (**B**), CR-1-31-B (**C**), and zotatifin (**D**), and after 24 hours of incubation cell viability was assessed using the MTT method after 24 hours of incubation. The data represent the mean ± standard deviation of three independent experiments, each with four replicates. Statistical analysis was performed using a one-way ANOVA followed by a Dunnett’s post hoc test. ns, not significant; *, *p* < 0.05; **, *p* < 0.01; ****, *p* < 0.0001.

We then decided to evaluate the antiviral potential of zotatifin, relative to other rocaglates, against the Mayaro virus (MAYV), an emerging alphavirus in the *Togaviridae* family that poses a potential threat to the Americas (Wei et al., 2024). HDFs or HMC3 cells were pretreated with the indicated concentrations of rocaglates for 2 hours before being infected with MAYV at a MOI of 1. Viral titers in the rocaglate-treated cells’ supernatants were evaluated 24 hours post-infection using plaque-forming assays. Rocaglamide A treatment did not significantly reduce MAYV replication at non-toxic concentrations (Figure 2A). However, CR-1-31-B inhibited MAYV replication by 2-3 logs in both HDFs and HMC3 cells (Figure 2B and D). Zotatifin demonstrated potent inhibition of MAYV replication in both cell lines, achieving a greater than 4-log reduction in viral titers at 50 nM (Figure 2C and E).

**Figure 2.**
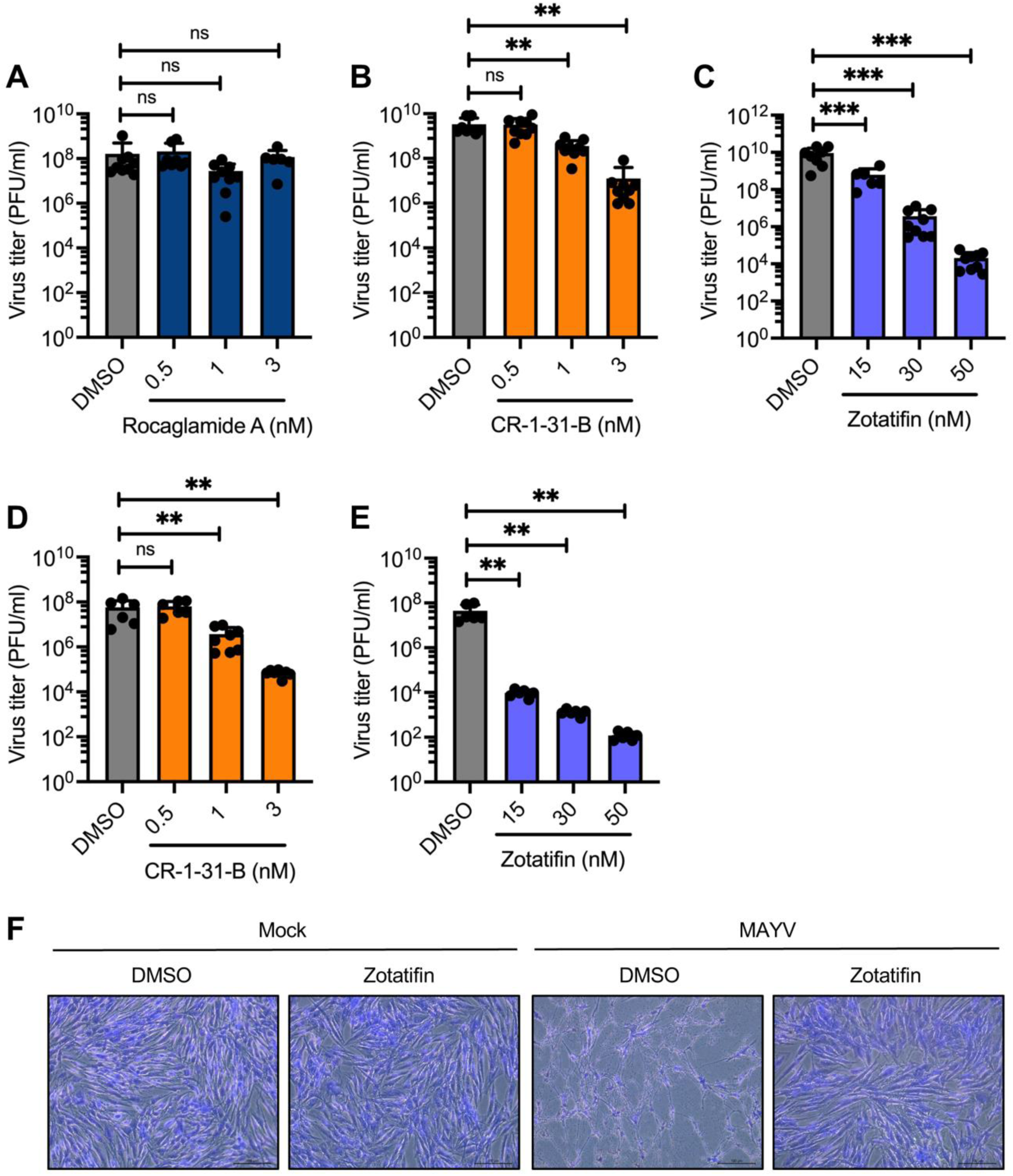
Zotatifin inhibits MAYV replication in a dose-dependent manner. HDFs (**A-C**) or HMC3 cells (**D-E**) were pretreated with the rocaglates at the indicated concentrations for two hours and then infected with the AVR0565 strain of MAYV at a MOI of 1. After 24 hours of infection in presence of the compounds, viral progeny production was evaluated using a plaque-forming assay. The data represent the mean ± standard deviation of three independent experiments in triplicate. Statistical analysis was performed using a one-way ANOVA, followed by a Dunnett’s post hoc test. ns, not significant; **, *p* < 0.01; and ***, *p* < 0.001. (**F**) HDFs were pretreated with zotatifin (50 nM), as described above, and then infected with MAYV (AVR0565 strain). After 48 hours of infection in presence or absence of zotatifin, the cells were fixed with a 4% formaldehyde solution and stained with a 2% crystal violet solution. Cell morphology was then evaluated using an inverted microscope. Scale bar: 100 μm.

As with other alphaviruses, MAYV induces robust cytopathic effects in several cell lines, including HDFs (Sugasti-Salazar et al., 2021). Microscopic examination at 48 hours post-infection revealed that zotatifin treatment completely protected HDFs against MAYV-induced cytopathic effects (Figure 2F), maintaining cell morphology like that of uninfected controls. Overall, these data demonstrate that zotatifin effectively controls MAYV replication in both HDFs and HMC3 cells.

### 3.2 Zotatifin reduces the expression of MAYV E1 and nsP1 proteins

Zotatifin is a potent inhibitor of the RNA helicase eIF4A, which plays a critical role in protein translation initiation (Ernst et al., 2020). Therefore, we analyzed the synthesis of MAYV E1 and nsP1 proteins following zotatifin treatment. Western blot analysis revealed that zotatifin treatment reduced the synthesis of both viral proteins in a dose-dependent manner in both HDFs (Figure 3A) and A549 cells (Figure 3B). Similar results were detected using immunofluorescence assays. We observed a reduction in the levels of MAYV E1 and nsP1 proteins in the A549 cells treated with 50 nM zotatifin (Figure 3C), a concentration that was well tolerated by the cells (Figure 3D). These results indicated that zotatifin significantly downregulates MAYV E1 and nsP1 protein expression.

**Figure 3.**
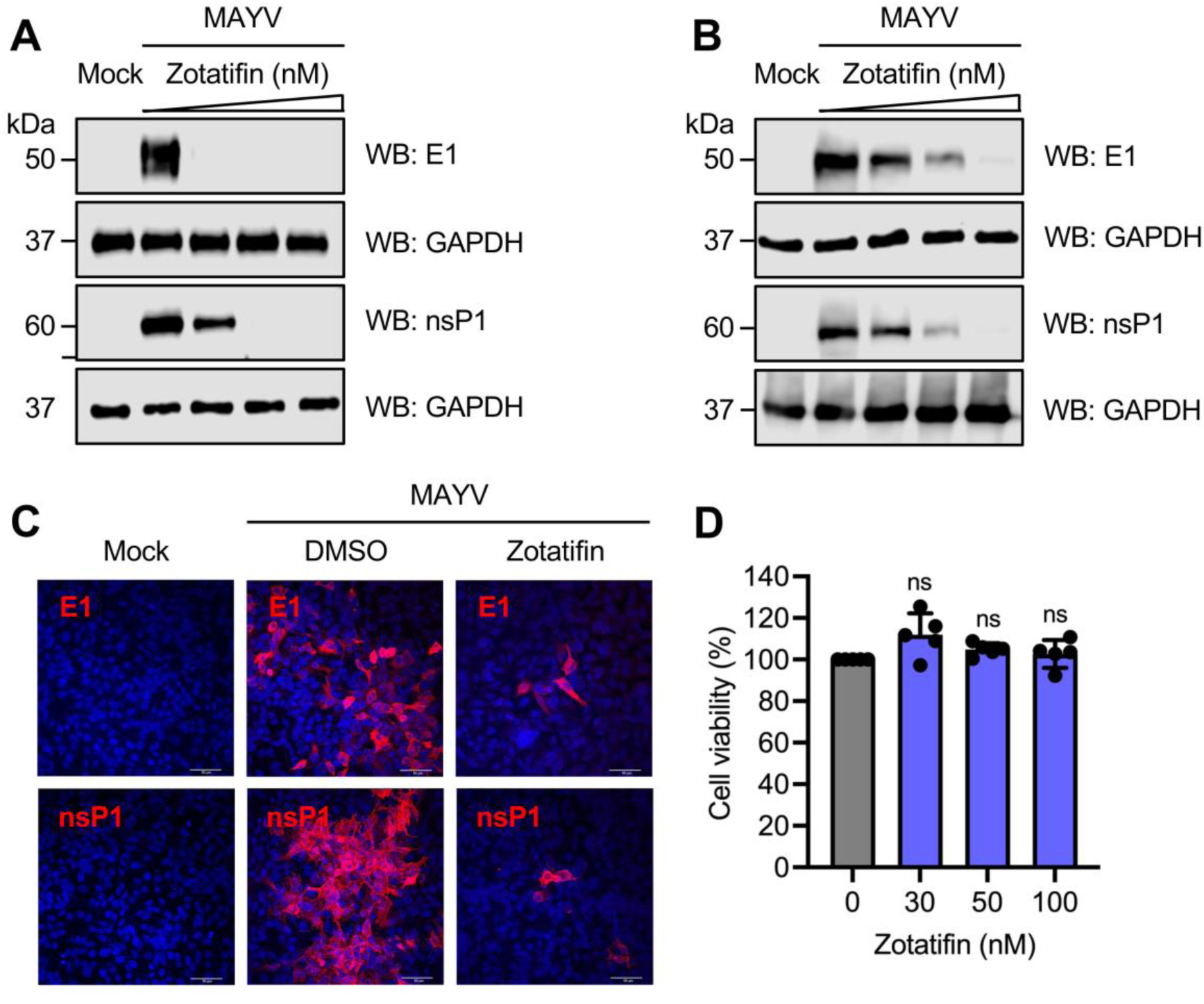
Zotatifin reduces MAYV E1 and nsP1 protein levels. HDFs (**A**) or A549 cells (**B**) were pretreated with zotatifin at doses of 15, 30, and 50 nM. Cells were then infected with MAYV AVR0565 strain and treated with the compound, as previously described. At 24 hours after infection, the levels of the E1 and nsP1 viral proteins were evaluated using Western blot analysis. One representative image of three independent assays is shown. Glyceroaldehyde-3-phosphate dehydrogenase (GAPDH) protein was used as a loading control. (**C**) A549 cells were treated or not with 50 nM zotatifin and then infected with MAYV. After 24 hours, MAYV E1 and nsP1 proteins were analyzed using an immunofluorescence assay. One representative microphotograph of at least 10 fields is shown. Scale bar: 50 μm. (**D**) A549 cells were treated with the indicated doses of zotatifin. After 48 hours of incubation, cell viability was evaluated using the MTT method. The data represent the mean ± standard deviation of one experiment with five replicates. Statistical analysis was carried out using a one-way ANOVA, followed by a Dunnett’s post hoc test. ns, not significant.

### 3.3 Zotatifin exhibits strain-independent antiviral activity against MAYV

We then decided to study zotatifin’s ability to inhibit the replication of different MAYV strains isolated from various regions in Latin America. We treated HDFs with or without 50 nM zotatifin for 2 hours and then infected with MAYV strains from Trinidad and Tobago (TRVL 4675), Brazil (BeH256), or Suriname (D218). Zotatifin significantly reduced viral titers by approximately 4-5 logs for all tested MAYV strains (Figure 4A-C), indicating strain-independent antiviral efficacy.

**Figure 4.**
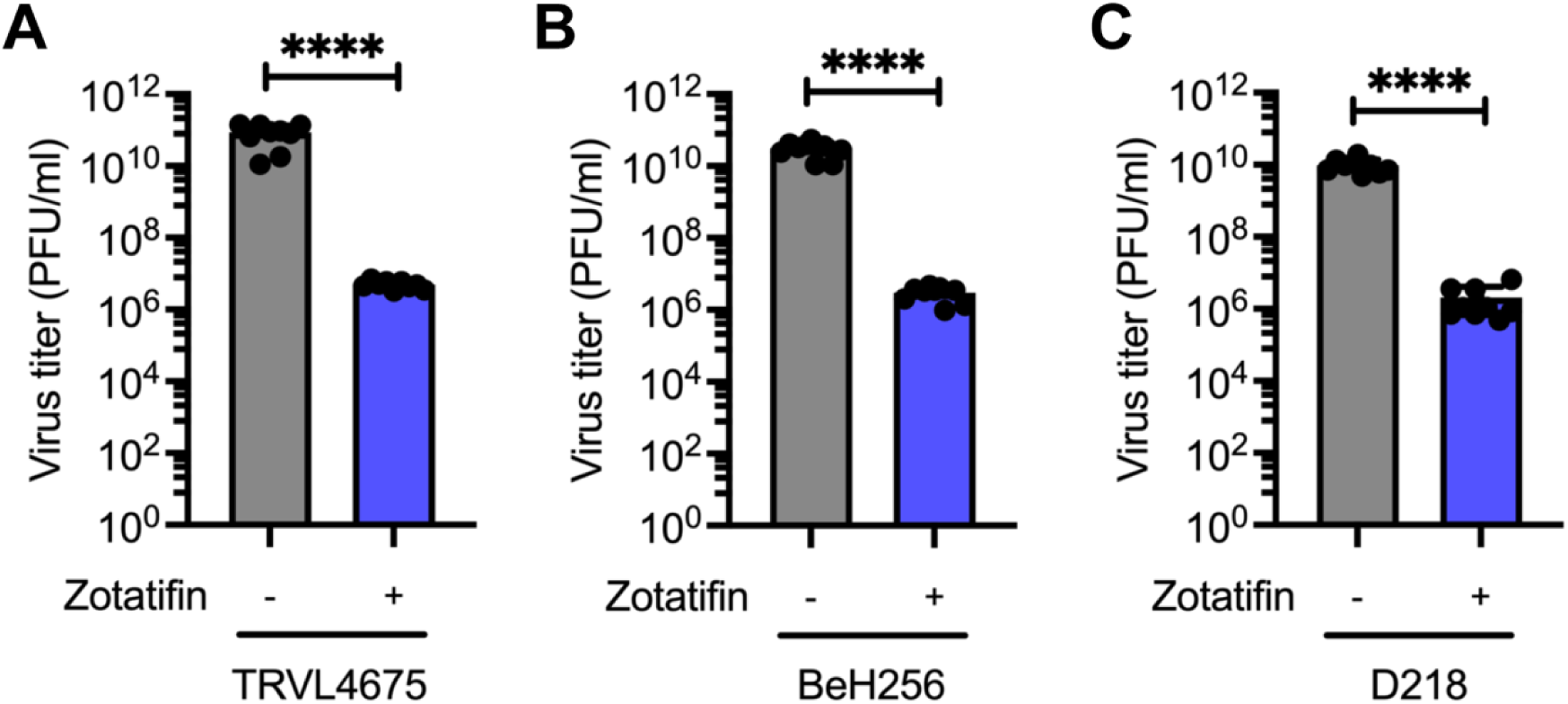
Zotatifin inhibits MAYV replication in a strain-independent manner. HDFs were pretreated with zotatifin (50 nM) and then infected with the following MAYV strains: TRVL4675 (Trinidad and Tobago) (**A**), BeH256 (Brazil) (**B**), or D218 (Surinam) (**C**). After 24 hours of infection in presence or absence of the compound, viral progeny production was assessed using a plaque-forming assay. The data represent the mean ± standard deviation of three independent experiments in triplicate. Data were analyzed using an unpaired t-test. ****, *p* < 0.0001.

### 3.4 Zotatifin acts through several mechanisms in the MAYV life cycle

Time-of-addition studies were conducted to determine zotatifin’s mechanism of action against MAYV (Figure 5A). Pretreatment of HDFs with zotatifin for only 2 hours prior to infection reduced MAYV titers by approximately 2 logs (Figure 5B, *p* < 0.01), suggesting the activation of cellular antiviral pathways by the compound. During the binding assay, in which zotatifin was only present during virus attachment at 4°C, no significant reduction in viral titers was observed (Figure 5C), indicating that zotatifin does not interfere with virus binding. Similarly, the entry assay showed no significant effect when zotatifin was present only during the 2-hour entry period (Figure 5D). However, adding zotatifin 2 hours post-infection and maintaining it for 24 hours resulted in a significant reduction in viral titer (Figure 5E, *p* < 0.001), which is consistent with the inhibition of viral protein synthesis through eIF4A targeting. These results suggest that zotatifin may act through several mechanisms: activation of cellular antiviral responses during pretreatment and direct inhibition of viral protein synthesis during the post-entry phase.

**Figure 5.**
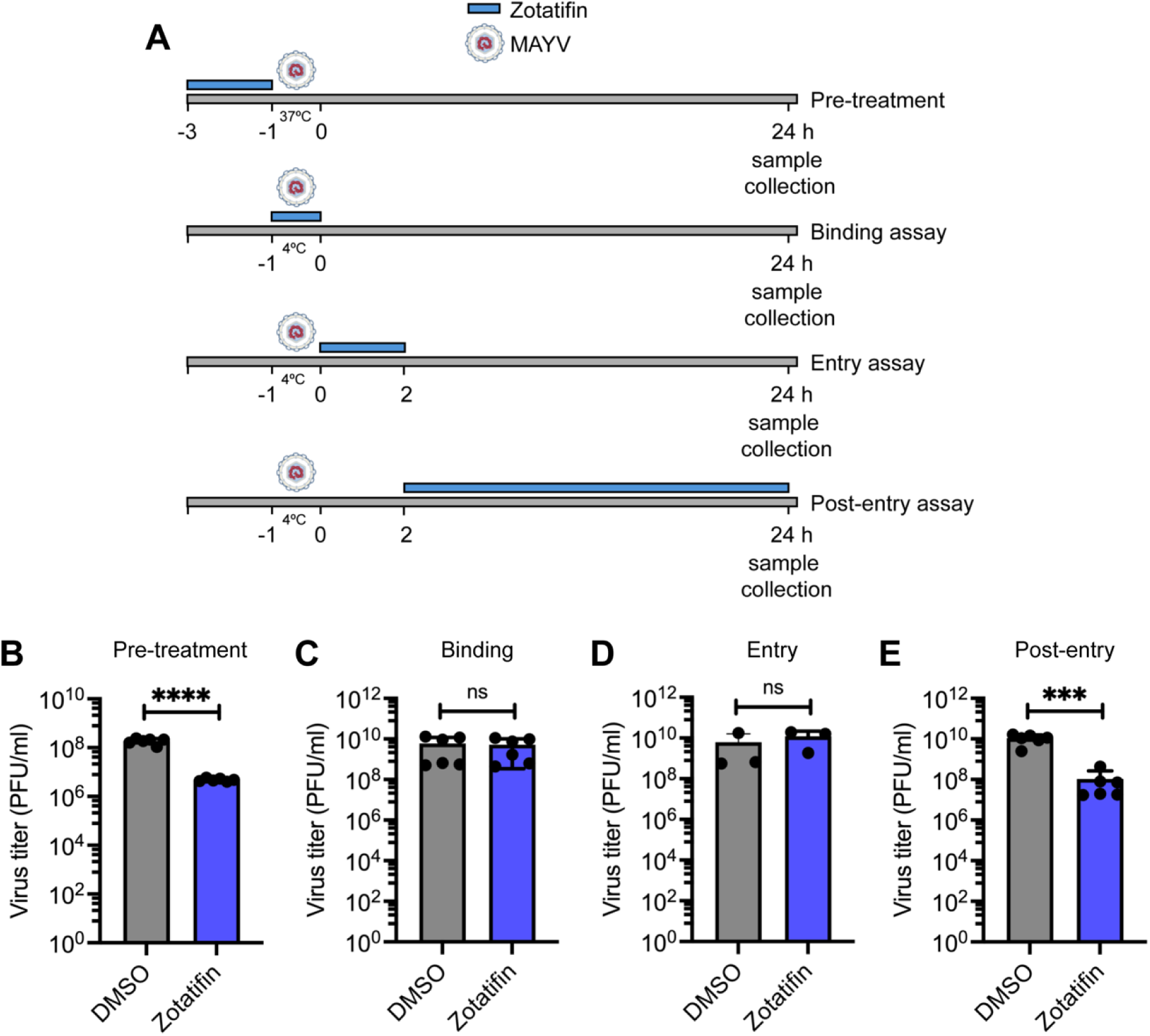
Zotatifin blocks MAYV replication when cells are pretreated or during late stages of the viral cycle. (**A**) Diagram showing the pre-treatment, binding, entry, and post-entry assays performed. Viral quantification in the supernatants of HDFs treated or untreated with zotatifin and infected with MAYV (AVR0565 strain) as indicated in A: in pre-treatment (B), binding (**C**), entry (**D**), and post-entry (**E**) assays. Data represent the mean ± standard deviation of two independent experiments in triplicate. Data were analyzed using an unpaired t-test. ns, not significant; **, *p* < 0.01; and ***, *p* < 0.001.

### 3.5 Zotatifin activates type I interferon signaling pathways in A549 cells

The antiviral effect observed with short-term zotatifin pretreatment, combined with previous reports of zotatifin-induced interferon-stimulated genes (ISGs) (Zhao et al., 2023), prompted us to investigate the potential activation of the IFN pathway in A549 cells. RT-qPCR analysis of A549 cells treated with 50 nM zotatifin for 8 hours revealed significant upregulation of *IFNα* (2.49-fold, *p* < 0.001), *DDX58* (2.19-fold, *p* < 0.0001), *MxA* (6.23-fold, *p* < 0.001), and *ISG15* (8.21-fold, *p* < 0.01) compared to DMSO-treated controls (Figure 6). These results indicate that zotatifin can activate type I IFN signaling, providing a mechanistic basis for its antiviral efficacy when used as a pretreatment.

**Figure 6.**
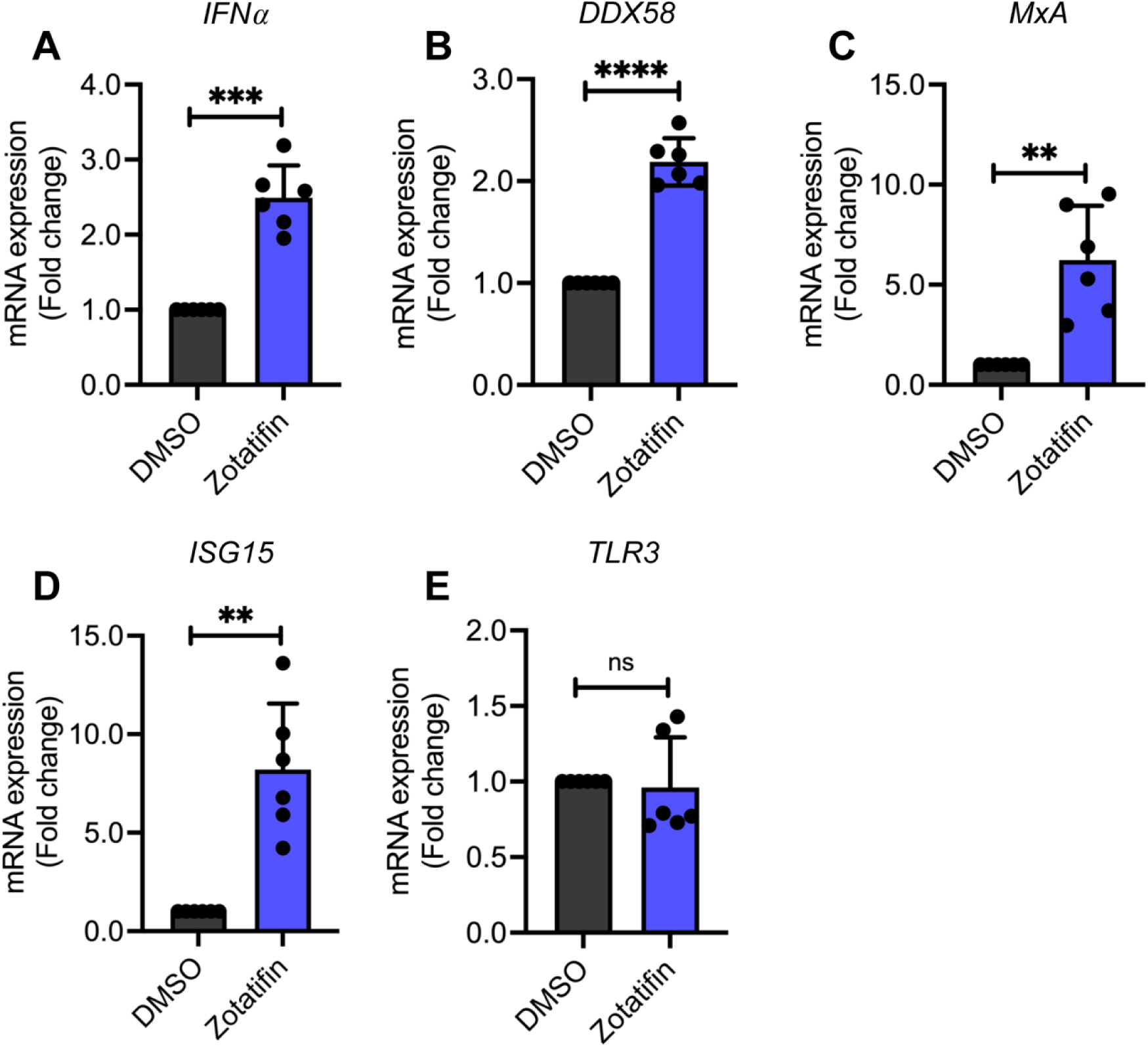
Zotatifin upregulates interferon and interferon-stimulated genes in A549 cells. A549 cells were treated or untreated with 50 nM zotatifin for 8 hours. Next, the relative transcription levels of *IFNα* (**A**), *DDX58* (**B**), *MxA* (**C**), *ISG15* (**D**), and *TLR3* (**E**) genes were analyzed using RT-qPCR. Relative mRNA expression was calculated using the β-actin gene for normalization with the ΔΔCT method. Data represent the mean ± standard deviation of two independent experiments in triplicate. Data were analyzed using an unpaired t-test. ns, not significant; **, *p* < 0.01; ***, *p* < 0.001; and ****, *p* < 0.0001.

### 3.6 Zotatifin demonstrates broad-spectrum activity against arboviruses

To evaluate the antiviral breadth of zotatifin, we tested its efficacy against several medically important arboviruses: Chikungunya (CHIKV), Una (UNAV), and Zika (ZIKV). HDFs were treated with zotatifin and then infected with each virus at an MOI of 1. Zotatifin significantly reduced viral titers in a dose-dependent manner for CHIKV (4-log reduction at 50 nM, *p* < 0.0001), UNAV (>2 log at 50 nM, *p* < 0.01), and ZIKV (>1-log reduction at 50 nM, *p* < 0.05) (Figure 7A, B, and C).

**Figure 7.**
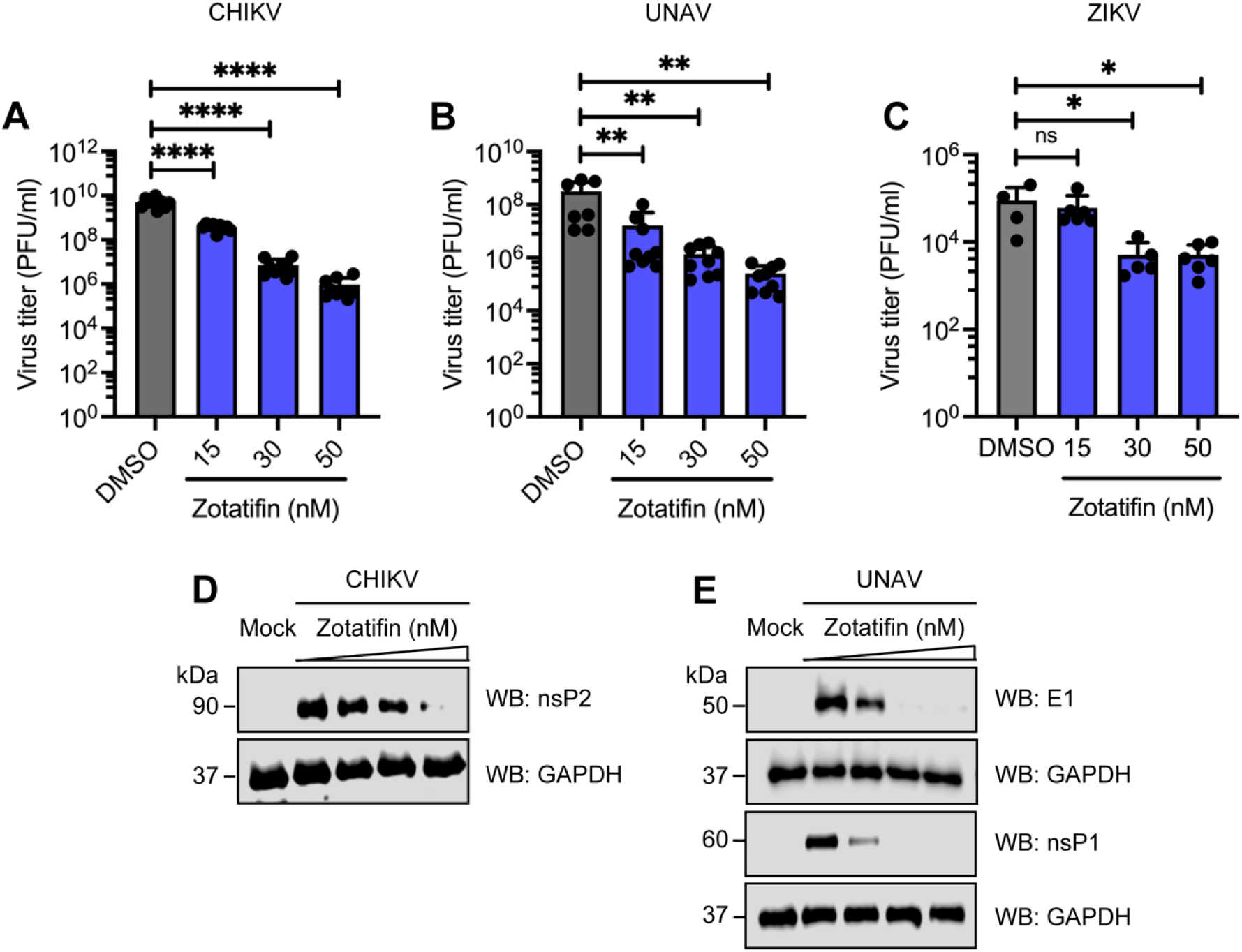
Zotatifin inhibits the replication of the arbovirus Chikungunya, Una, and Zika, in a dose-dependent manner. HDFs were pretreated with zotatifin at the indicated concentrations and then infected with CHIKV (**A**), UNAV (**B**), or ZIKV (**C**), as previously described. After 24 hours of infection, viral progeny production in the cell culture supernatants was quantified using a plaque-forming assay. Data represent the mean ± standard deviation of three independent experiments in triplicate. Statistical analysis was carried out using a one-way ANOVA followed by a Dunnett’s post hoc test. *, *p* < 0.05; **, *p* < 0.01; ****, *p* < 0.0001. HDFs were pretreated with Zotatifin, then infected with CHIKV or UNAV. Next, the levels of the viral nsP2 (**D**), E1, and nsP1 (**E**) proteins were analyzed using Western blot assays. One representative image of three independent assays is shown. Glyceroaldehyde-3-phosphate dehydrogenase (GAPDH) protein was used as a loading control.

Finally, we evaluated whether zotatifin treatment affected the synthesis of viral proteins. We observed a reduction in CHIKV nsP2 and UNAV E1, and nsP1 protein levels in response to zotatifin in a dose-dependent manner (Figure 7D and E). These results demonstrate that zotatifin exhibits broad-spectrum antiviral activity against multiple arbovirus members.

### 3.7 Zotatifin inhibits viruses from *Orthomyxoviridae*, *Rhabdoviridae* and *Poxviridae* families

To further evaluate zotatifin’s broad-spectrum potential, we assessed its efficacy against three viruses: influenza A (*Orthomyxoviridae* family), vesicular stomatitis (*Rhabdoviridae* family), and vaccinia (*Poxviridae* family). A549 cells were pretreated with or without zotatifin, then infected with a recombinant influenza virus or recombinant vesicular stomatitis carrying a GFP reporter gene (PR8-GFP and rVSV-GFP, respectively). At 24 hours after infection, we evaluated the GFP intensity using flow cytometry. Zotatifin treatment significantly decreased GFP intensity, regardless of the virus or MOI tested (Figure 8A, B, E, and F), indicating that zotatifin inhibits the replication of both viruses. Additionally, we quantified the virus titers in the cell culture supernatant from treated and untreated cells PR8-GFP and rVSV-GFP infected cells, and in the lysates from treated and untreated VV infected cells. We observed a significant reduction in PR8-GFP and rVSV-GFP titers (Figure 8C, D, and G). A similar reduction in viral titer was also observed in VV infected cells upon treatment with zotatifin (Figure 8H-I).

**Figure 8.**
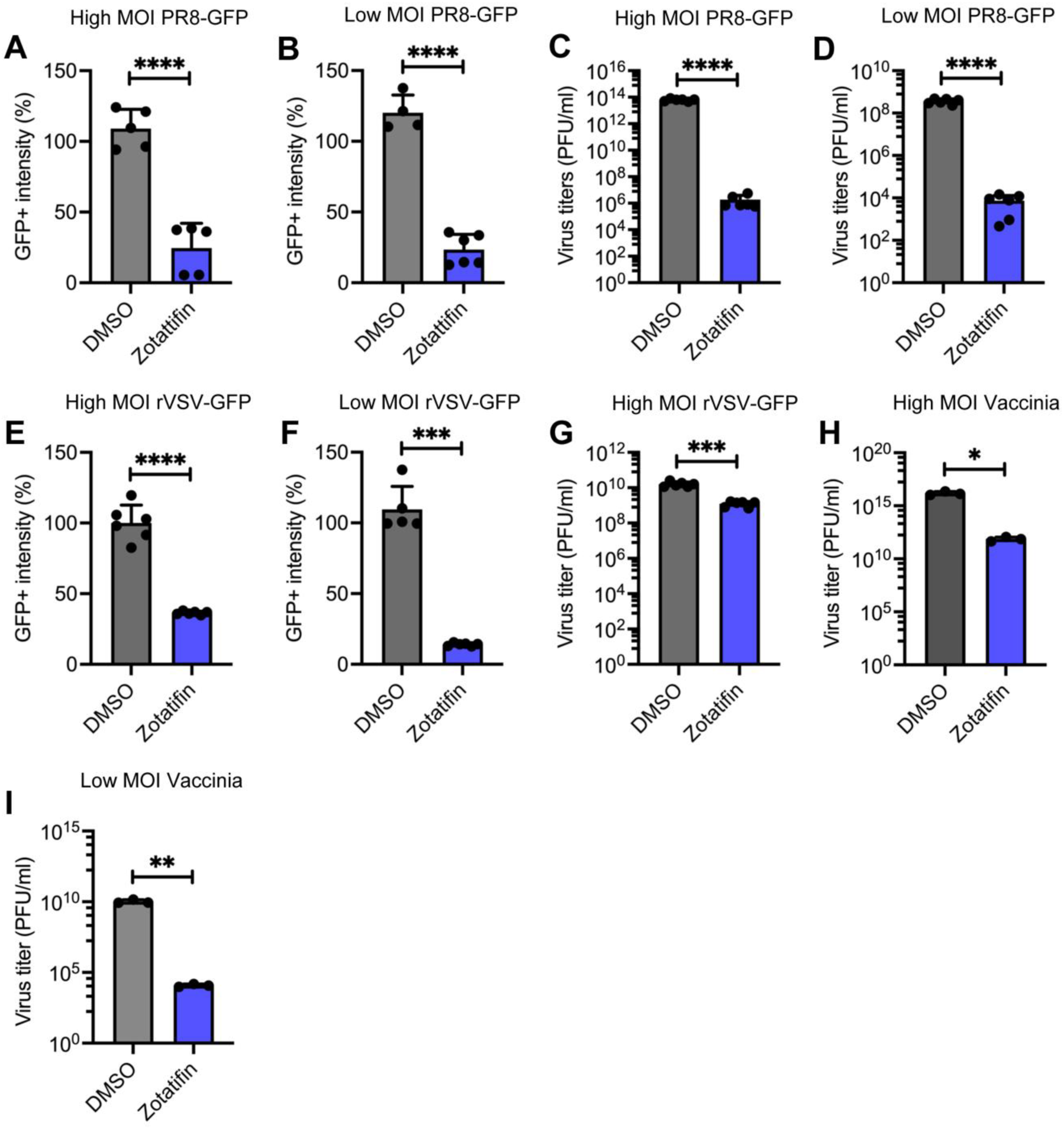
Zotatifin inhibits the replication of influenza A, vesicular stomatitis, and vaccinia viruses. A549 cells were pretreated with zotatifin, then infected with PR8-GFP (**A-D**), rVSV-GFP (**E-G**) or vaccinia (**H-I**) viruses at low (0.5) or high (5) multiplicity of infection (MOI). After 24 hours, the intensity of GFP, and the quantity of viral progeny in PR8-GFP and rVSV-GFP infected cells were assessed using flow cytometry, and a plaque-forming assay. Virus titers in the cell lysates of vaccinia virus infected cells were determined by plaque assay. The data represent the mean ± standard deviation of two independent experiments in triplicate. Statistical analysis was carried out using a one-way ANOVA followed by a Dunnett’s post hoc test or an unpaired t-test. ns, not significant; *, *p* < 0.05; **, *p* < 0.01; ***, *p* < 0.001; and ****, *p* < 0.0001.

Finally, we analyzed the effect of zotatifin on viral protein synthesis. We observed a reduction in influenza A, VSV, and vaccinia protein levels in those cells treated with zotatifin, regardless of the MOI tested (Figure 9 A-F). These findings suggest that zotatifin exhibits an antiviral activity against IAV, VSV, and vaccinia viruses.

**Figure 9.**
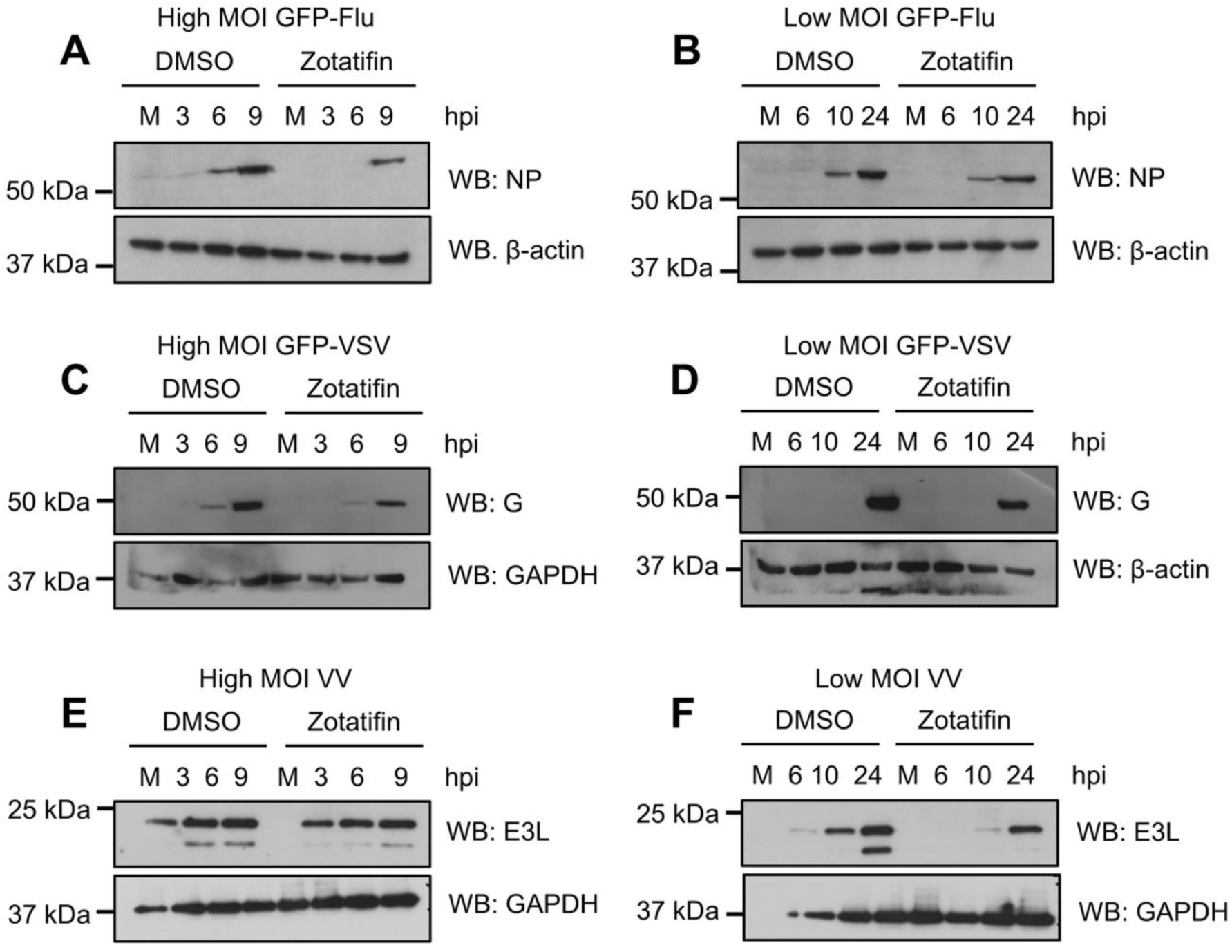
Zotatifin downregulates the synthesis of viral proteins in influenza A virus, vesicular stomatitis virus, and vaccinia virus infected cells. A549 cells were pretreated with zotatifin, then infected with PR8-GFP (**A-B**), rVSV-GFP (**C-D**), or vaccinia (**E-F**) viruses, as previously indicated. At the indicated hours after infection, levels of influenza A NP, VSV M, or vaccinia E3L proteins were evaluated using Western blot analysis. One representative image of three independent assays is shown. β-actin or GAPDH proteins were used as a loading controls.

## 4. Discussion

The inhibition of eIF4A by the natural rocaglate silvestrol has been reported to exhibit antiviral activity against the ZIKV, human rhinovirus, coronavirus, EBOV, and picornavirus (Biedenkopf et al., 2017; Elgner et al., 2018; Muller et al., 2018). Zotatifin, a synthetic rocaglate, exhibits significant antiviral activity against various coronaviruses, including HCoV-229E, MERS-CoV, and SARS-CoV-2 (Ernst et al., 2020; Gordon et al., 2020; Obermann et al., 2022). However, zotatifin’s potential for broad-spectrum antiviral activity remains incompletely characterized.

In this study, we evaluated zotatifin’s potential to control the replication of viruses from five families: *Orthomyxoviridae*, *Rhabdoviridae*, *Flaviviridae*, *Togaviridae*, and *Poxviridae*. Our results revealed that, compared with the rocaglates rocaglamide A and CR1, zotatifin exhibits reduced cytotoxicity on HDF and HMC3, which is consistent with the reduced cytotoxic effects on immune cells previously reported (Obermann et al., 2022). We also show that zotatifin potently reduces both MAYV replication and viral protein synthesis in a dose-dependent manner, regardless of the cell line or MAYV strain tested. Additionally, we demonstrated that zotatifin effectively controls the replication of other togaviruses, including CHIKV and UNAV, and inhibits the medically relevant ZIKV flavivirus. These findings suggest that zotatifin exhibits antiviral activity against diverse arboviruses with similar epidemiological characteristics, thus expanding the therapeutic potential of this compound. Importantly, we demonstrated that zotatifin also downregulates the replication and protein synthesis of other, unrelated viruses, such as influenza A virus, vesicular stomatitis virus, and vaccinia virus, confirming its broad-spectrum antiviral properties previously suggested (Taroncher-Oldenburg et al., 2021).

Time-of-addition studies revealed that adding zotatifin 2 hours post-infection effectively reduced MAYV replication, indicating a post-entry antiviral effect. This observation is consitent with the inhibition of protein translation by targeting the host helicase eIF4A (Ernst et al., 2020; Obermann et al., 2022). Additionally, we demonstrated that pretreating cells with zotatifin for just 2 hours provides antiviral protection against MAYV. Our data demonstrating that IFN and IFN-response genes are upregulated in A549 cells treated with zotatifin together with previous reports showing that zotatifin induces the IFN pathway (Zhao et al., 2023), led us to propose that zotatifin’s antiviral activity may involve both direct translation inhibition and innate immune activation. However, additional host pathways may also contribute to zotatifin’s antiviral effects (Chen et al., 2021; Ho et al., 2021).

Although zotatifin shows promise as an antiviral treatment, several limitations must be recognized. Our studies were performed exclusively in vitro, which may not fully recapitulate the complexity of viral infections in vivo. Moreover, the tested viruses represent only a subset of medically important viruses.

To our knowledge, this represents the first comprehensive demonstration of zotatifin’s broad-spectrum antiviral activity across multiple virus families. Zotatifin’s antiviral activity against diverse MAYV strains from different geographic locations, suggest that this compound may be effective against genetically variable viruses, an important factor to consider in the contex of emerging viruses. Moreover, zotatifin’s broad-spectrum antiviral activity and safety profile demonstrated in preclinical and clinical studies (Boyer et al., 2025; Gerson-Gurwitz et al., 2021; Kuzuoglu-Ozturk et al., 2025; Manara et al., 2025), supports its consideration as an antiviral agent for emerging and reemerging viral diseases.

## Credit authorship contribution statement

Conceptualization, P.V.-T., D.C., C.R., and J.G.-S.; methodology, P.V.-T., D.C., P.E.G.-J., D.Z., I.T.-L., F.G.-C., J.C.M., M.B., C.R., and J.G.-S.; Validation, P.V.-T., D.C., C.R., and J.G.-S.; formal analysis, P.V.-T., D.C., C.R., and J.G.-S.; investigation, P.V.-T., D.C., P.E.G.-J., D.Z., I.T.-L., F.G.-C., J.C.M.; M.B., C.R., and J.G.-S.; resources, C.R. and J.G.- S.; writing-original draft preparation, C.R. and J.G.-S.; writing-review and editing, P.V.-T., D.C., P.E.G.-J., D.Z., I.T.-L., F.G.-C., J.C.M.; M.B., C.R., and J.G.-S.; supervision, P.V.-T., D.C., C.R., and J.G.-S.; project administration, P.V.-T., C.R., and J.G.-S.; funding acquisition, P.V.-T., C.R., and J.G.-S. All authors have read and agreed to the published version of this manuscript.

## Acknowledgments

We would like to thank Scott Weaver (WRCEVA, UTMB, USA) for providing the Mayaro and Una virus strains; Adolfo García-Sastre for providing the influenza virus strain; and Carmen Rivas for providing the A549 cells. We are also grateful to Rodolfo Contreras and Nicanor Obaldía for their support with laboratory facilities. Finally, we thank Carolina Alvardo and Armando Castillo for their support and for granting us access to the confocal microscope at INDICASAT.

## Declaration of generative AI use

We used Scispace to search for information, review the manuscript, and make language corrections.

